# Improved methods for the large-scale cultivation of *Giardia lamblia* trophozoites in the academic laboratory

**DOI:** 10.1101/2022.05.12.491643

**Authors:** Brian T. Wimberly, Danielle Bilodeau, Olivia S. Rissland, Jeffrey S. Kieft

**Affiliations:** Department of Biochemistry & Molecular Genetics and the RNA Biosciences Initiative, University of Colorado Anschutz Medical Campus, School of Medicine, Aurora, CO 80045

## Abstract

The parasitic protist *Giardia lamblia* causes giardiasis, one of the leading diarrheal diseases worldwide. As a phylogenetically divergent eukaryote, *Giardia* is also an emerging model organism for exploring the diversity of core eukaryotic metabolic pathways and important molecular structures. To date, methods for the cultivation of large numbers (>10^9^) of *Giardia* trophozoites have been either expensive or cumbersome, hampering structural and biochemical studies. Here, we present two improved strategies for the laboratory-scale growth of billions of adherent trophozoites, with suggestions for the optimization of media components, media preparation, and harvesting methods.

## Introduction

Infection by the protist *Giardia lamblia* leads to giardiasis, the most common parasitic human diarrheal disease and a common cause of childhood diarrhea, especially in developing nations (Pires *et al*., 2015). Although giardiasis is typically not as life-threatening as diarrhea from some other pathogens (Kotloff *et al*., 2013), even asymptomatic chronic infections from *Giardia* have adverse stunting effects on development (Kirk *et al*., 2010, Rogawski *et al*., 2018). By infecting other mammals including pets and livestock, the parasite is also economically significant (Feng and Xiao, 2011). Disturbingly, the incidence of treatment-refractory giardiasis is increasing in some areas, making the discovery of novel therapies ever more important (Lalle and Hanevik, 2018). To find new treatments, a deeper understanding of *Giardia’s* biology is needed to identify targetable idiosyncrasies of this unusual pathogen.

As a highly divergent eukaryote, *Giardia* probably has many such targetable features. A microaerophilic and parasitic protist, it lacks canonical mitochondria, and many of its structures and metabolic pathways appear to be greatly streamlined compared to those from other eukaryotes (Morrison *et al*., 2007). Early ribosomal RNA-based analyses of its phylogenetic position suggested that *Giardia* and other closely related diplomonads may be a primitive lineage that branched off very early from other eukaryotes (Edlind and Chakraborty 1987). However, more recent work suggests that the diplomonads may have diverged much more recently, perhaps because of adoption of a parasitic lifestyle (*e.g*., Hashimoto *et al*., 1998). Whatever its causes, the highly divergent phylogenetic position of *Giardia* is of great interest to biologists seeking to define the limits of conservation of the core metabolic pathways and structures in eukaryotes. While its tetraploid genome is inconvenient for genetics, *Giardia* is nevertheless emerging as an important new model organism. Significantly, several strains of *Giardia* can be cultured axenically (*i.e*., in the absence of other organisms).

Many structural biology, biophysical, and biochemical methods require milligrams of a pure macromolecule, often obtained by overexpression in a heterologous expression system. However, many larger macromolecular assemblies such as ribosomes cannot be overexpressed in a foreign host and must be isolated from larger-scale cultures of the target species. When beginning to study the structure and function of the *Giardia* ribosome (Eiler *et al*., 2021), our early preparations using normal or only lightly modified cultivation methods yielded several hundred million trophozoites, from which only a few tens of micrograms of *Giardia* ribosomes were obtained. This yield is only about 1% of the ribosome yield from 100 mL rabbit reticulocyte lysate, a thoroughly studied eukaryotic model system. New larger-scale cultivation methods proved necessary to obtain sufficient material, even for the sample-efficient cryo-EM methods (Eiler *et al*., 2021).

In rapidly growing *Giardia* cultures, almost all trophozoites adhere to the walls of the culture vessel, with most of the dying and dead cells floating in media or precipitated at the bottom. Accordingly, it is best practice during harvest to discard cells in the media and collect only the adherent trophozoites. Most small-scale growths are performed in 10-15 mL flat-wall tubes with an unoptimized surface-to-volume ratio, meaning that the yield is limited by the available surface area rather than the ability of the media to support growth. Therefore, all strategies for improved larger-scale cultivation methods have included a way to increase the surface-to-volume ratio. One group designed their own double-sided or “inside-out” glass roller bottles that required custom fabrication (Farthing *et al*., 1982). Another early solution featured insertion of many open-ended glass cylinders into a 19-liter glass carboy, with a pump used for sterilization, cleaning, and to recirculate media (Wieder *et al*., 1983). More recently, and on a smaller scale, autoclavable polypropylene drinking straws were inserted into 500 mL glass media bottles (Paredez *et al*., 2014). Yet another recently reported strategy is to add glass beads to culture vessels (Serradell *et al*., 2016). While all these strategies improve the surface-to-volume ratio, each also suffers from disadvantages. The older methods depend on an elaborate custom-built apparatus; the more recent work is more convenient, but also of a smaller scale. Importantly, none of these strategies allows noninvasive monitoring of growth, a highly desirable goal for several reasons, such as variability in batch-to-batch media preparations. Here we present two low-cost and complementary strategies for the cultivation of billions of *Giardia* trophozoites.

## Methods and Results

### Choice and preparation of growth media

Standard methods for the culture of *Giardia* in modified TYI-S-33 media (Keister 1983) have been reviewed recently (Fink *et al*., 2020). The focus here will be on modifications for a larger scale of cultivation and optional modifications to support faster growth.

To prepare TYI-S-33 media, dissolve each ingredient (except for the bile and FBS) in the order given in Table 1 while stirring at medium speed. Allow each ingredient to dissolve completely before proceeding to the next ingredient. In addition to promoting *Giardia* growth, the optional arginine-HCl additive improves the solubility of some of the components that might otherwise be filtered out. Once all components are dissolved, adjust the pH to 7.1 with concentrated NaOH while stirring rapidly, and then adjust the volume to 890 mL by adding MilliQ water. Stir for an additional 20-30 minutes. Insufficient stirring at this step will lead to loss of nutrients during filtration.

**Table 1.** Diamond’s TYI-S-33 media as modified by Keister (1983)

Optional ingredients are listed in parentheses (see media notes, below). For 1 liter.
|  |  |
| --- | --- |
| MilliQ water | ~800 mL |
| 4.38 mM K<sub>2</sub>HPO<sub>4</sub>·3H<sub>2</sub>O | 1.0 g |
| 4.41 mM KHPO<sub>4</sub> | 0.60 g |
| 34.2 mM NaCl | 2.0 g |
| Biosate peptone<sup>1</sup> | 30.0 g |
| (10-20 mM arginine-HCl) | (2.11-4.22 g) |
| 55.6 mM glucose<sup>2</sup> | 10 g |
| 11.4 mM cysteine-HCl-H<sub>2</sub>O | 2.0 g |
| 0.071 mM ferric ammonium citrate<sup>3</sup> | 1.0 mL @ 22.8 mg/mL |
| 1.14 mM ascorbic acid | 0.20 g |
| 0.5-0.8 mg/mL bile extract<sup>4</sup> | 10-16 mL @ 50mg/mL |
| 10% v/v heat-inactivated fetal bovine serum<sup>5</sup> | 100 mL |

To sterilize, filter the media using a 0.45 micron filter in a biosafety cabinet, and then add 100 mL heat-inactivated FBS, 10-15 mL bile extract (from a 50 mg/mL sterile-filtered stock), and either 10 mL Pen/Strep (Gibco 15140-122) or 10 mL antibiotic-antimycotic (Gibco 15240-062).

The media performs best if used just after preparation, *i.e*., without long-term storage or freezing. Upon storage at 4°C, a pale precipitate containing in part cystine crystals (oxidized cysteine) will eventually form. Discard any media with a visible precipitate. If the media is not to be used the same day, aliquot into 50 mL Falcon tubes and freeze at −20°C.

### Inoculum

Small-scale starter cultures are initiated from frozen trophozoites as described elsewhere (Fink *et al*., 2020). These small-scale cultures are grown at 37°C in Nunclon delta flat-sided tubes (NUNC CS450), each tube containing about 11 mL of the media described above. After 48-72 h of growth, adherent cells should reach about 70-80% confluence (as observed with a tissue culture microscope). Cells should be passaged by inoculating 10-20 microliters of culture into another tube of fresh media. When starting from frozen cells, several passages may be necessary before consistent and fast growth is observed; doubling times for WB strains should be between 6 and 12 hours depending on media. When inoculating a culture of any size, the vessel should be filled so as to leave minimal air inside. Variation in the amount of residual air in the vessel can lead to variation in the length of the lag phase before exponential growth begins. In addition, cultures should be grown to no more than 70-80% confluence, as an overgrown inoculum will likely result in poor growth of subsequent large-scale culture. Once this 70-80% level of confluence is reached, the cells are harvested as described below and used to inoculate larger-scale cultures.

### General principles for harvesting G. lamblia

To harvest cells, first aspirate the media containing mostly defective and dead trophozoites, leaving the adherent cells. Two methods to detach the cells are (1) using decreased temperature, and (2) using a natural product (formononetin). (1) Cells are traditionally detached by adding ice-cold isotonic detaching buffer (*e.g*., PBS), followed by immersion of the vessel in ice water. A twenty-minute incubation in an ice/water slurry is sufficient. Tapping and/or inversion helps dislodge the cells. (2) Alternatively, the addition of 10 micromolar formononetin may be used to detach instead of lowering the temperature. This natural product disrupts the trophozoite cytoskeleton and flagella, resulting in detachment in as few as five minutes (Lauwaet *et al*., 2010). Detachment efficiency is consistently 10-20% higher with formononetin than with chilling. However, while formononetin does not kill *Giardia* trophozoites, it is possible that the compound may have undesirable or uncharacterized effects on metabolism.

With either method, both cysteine-HCl (to 1 mg/mL) and ascorbate (to 0.2 mg/mL) should be added to the detaching buffer, since these reducing agents have a synergistic effect on improving viability when trophozoites are exposed to oxygen (Gillin and Diamond 1981). Once detachment is complete, detached cells are pelleted at 1500xg for 5 minutes and then resuspended in a convenient volume of fresh buffer or fresh media. If inoculating a larger-scale culture, enough inoculum is added to yield a final concentration of 3,000-5,000 trophozoites/mL.

Below, we present two novel set-ups for large-scale growth, based on the goal of increasing the surface-to-volume ratio.

### The beehive: a new high-surface area system for large-scale cultivation

The first method uses a commercially available (Globe Scientific), autoclavable polyoxymethylene slide rack that fits snugly into a rectangular slide staining dish (Fig. 1; see Table 2 for components and part numbers). We name this setup the “beehive” based on its similarity to the boxes used to cultivate bee colonies. The system is compact and more space-efficient than an equivalent commercially available anaerobic system, which is designed to contain up to ~20 10 cm plates (the BD GasPak system) (Fig. 1A). Once loaded with glass slides, racks are inserted into staining dishes and autoclaved, preferably with a polypropylene tray for secondary containment (Fig. 1D). Once in a biosafety cabinet, the dishes are filled with inoculated media. Four such rack and dish assemblies require about 650-700 mL of media. Each dish is sealed with its lid, and then placed in an airtight vessel for secondary containment (a 5 liter Kimchi fermentation container works well), along with three BD GasPak carbon dioxide sachets to minimize contamination by oxygen.

**Figure 1.**
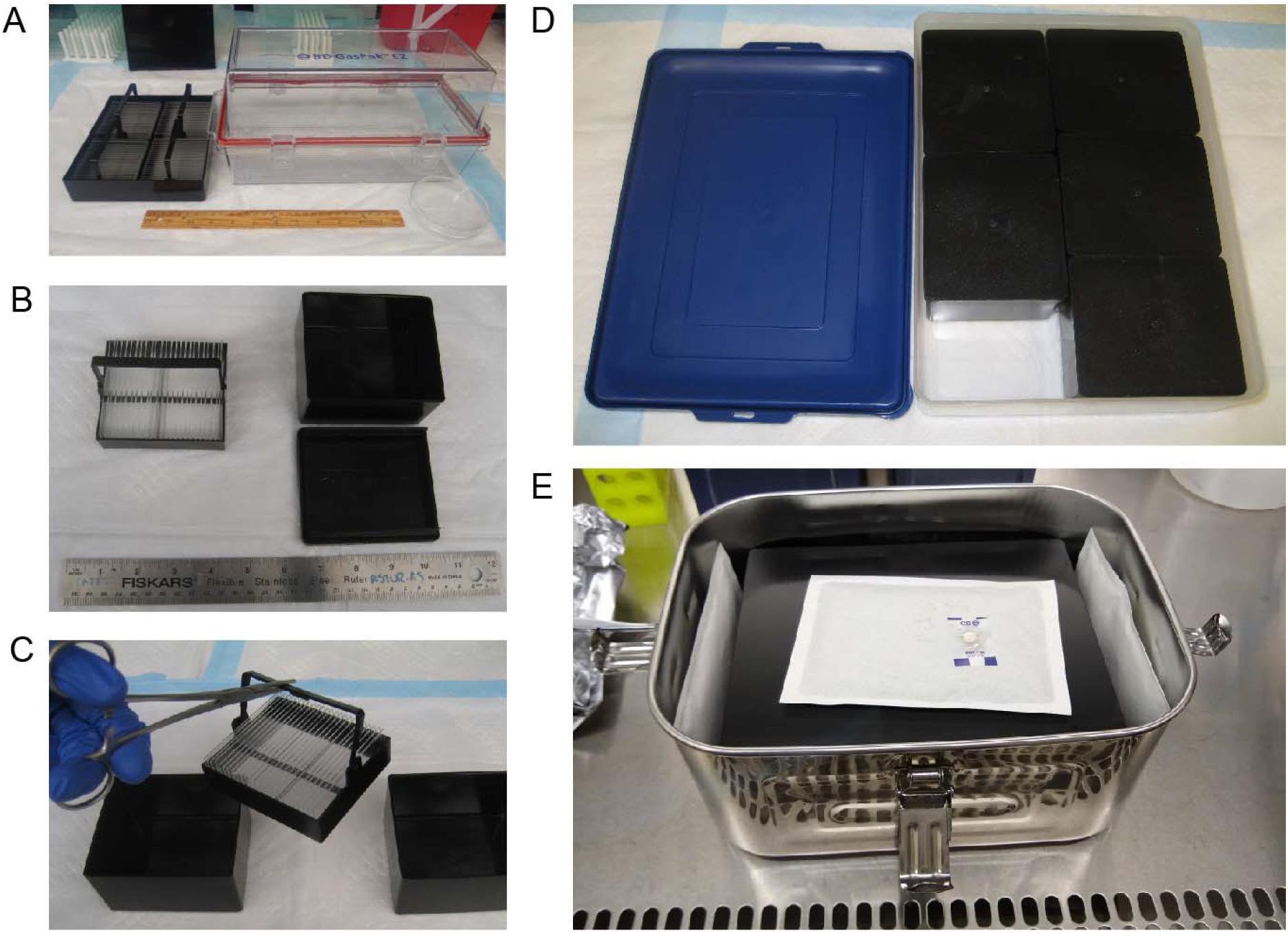
The ‘beehive’ system. **A.** Comparison of the dimensions of 4 beehive racks (which contain ~700 mL media and ~4000 cm^2^ area) and one BD GasPak container (~180 mL media and ~990 cm^2^ area in 18×10 cm plates). **B.** Top view of one loaded 25-slide rack and staining dish. **C.** Handling racks with hemostat. **D.** Loading racks and dishes into a polypropylene tray for autoclaving. **E.** Airtight Kimchi container ready to seal. Three BD GasPaks are used to reduce the oxygen concentration.

**Table 2.**
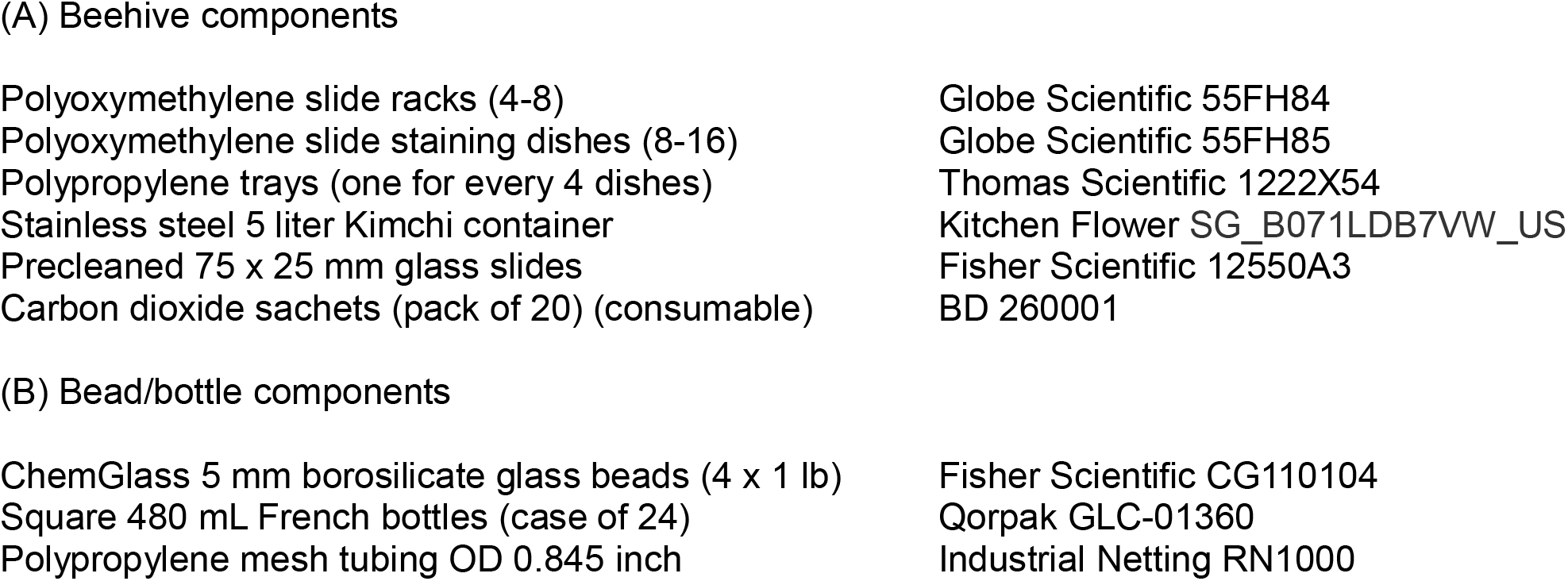
Composition of the beehive and bead/bottle systems.

**(A) Beehive components**
|  |  |
| --- | --- |
| Polyoxymethylene slide racks (4-8) | Globe Scientific 55FH84 |
| Polyoxymethylene slide staining dishes (8-16) | Globe Scientific 55FH85 |
| Polypropylene trays (one for every 4 dishes) | Thomas Scientific 1222X54 |
| Stainless steel 5 liter Kimchi container | Kitchen Flower SG_B071LDB7VW_US |
| Precleaned 75 x 25 mm glass slides | Fisher Scientific 12550A3 |
| Carbon dioxide sachets (pack of 20) (consumable) | BD 260001 |
| **(B) Bead/bottle components** |  |
| ChemGlass 5 mm borosilicate glass beads (4 x 1 lb) | Fisher Scientific CG110104 |
| Square 480 mL French bottles (case of 24) | Qorpak GLC-01360 |
| Polypropylene mesh tubing OD 0.845 inch | Industrial Netting RN1000 |

To harvest cells from the beehive, the containers are opened and then spent media is aspirated. Each rack is then lifted out of its dish and placed into a second pre-chilled dish containing detaching buffer, either chilled media or media containing formononetin. The detached cells are then harvested by centrifugation for 5 minutes at 1500xg. The late-log yield per dish with normal media is typically about 150 million cells and can be somewhat improved by scraping the dish used for growth.

The surface-to-volume ratio achieved by the beehive is higher than that seen by filling a 10 cm plate with 10 mL of media using the GasPak system. An interesting feature of the beehive system is that trophozoites can be transferred very rapidly (in a few seconds) from optimal growth conditions to a drastically new condition. One downside of the system is that it consists of literal black boxes, so the progress of growth cannot be monitored after inoculation. Another possible disadvantage is that the surface-to-volume ratio cannot be changed very much.

### The bead/bottle: a second new high-surface area system for large-scale cultivation

The second method we present is the “bead/bottle” (Table 2B and Fig. 2), an elaboration of the cultivation strategy described by Serradell *et al*. (2016) that uses a culture vessel filled with 5 mm diameter glass beads. Significantly, however, we made two improvements to improve function: (1) the use of a square, wide-mouth, almost flat-sided French glass bottle, and (2) the addition of one or more “windows,” each comprising a rigid polypropylene mesh tube that is placed in the bottom of the bottle. The tube excludes beads, and the flat side of the bottle allows undistorted visualization of adherent trophozoites using a microscope. Thus, together these changes allow noninvasive monitoring of trophozoite growth, if a tissue culture microscope with sufficient clearance is available (about 65 mm clearance is needed for the 480 mL bottle from QorPak). The wide mouth is necessary for easy insertion and positioning of the polypropylene tube, and it also facilitates filling with about 450 g of 5 mm diameter beads.

**Figure 2.**
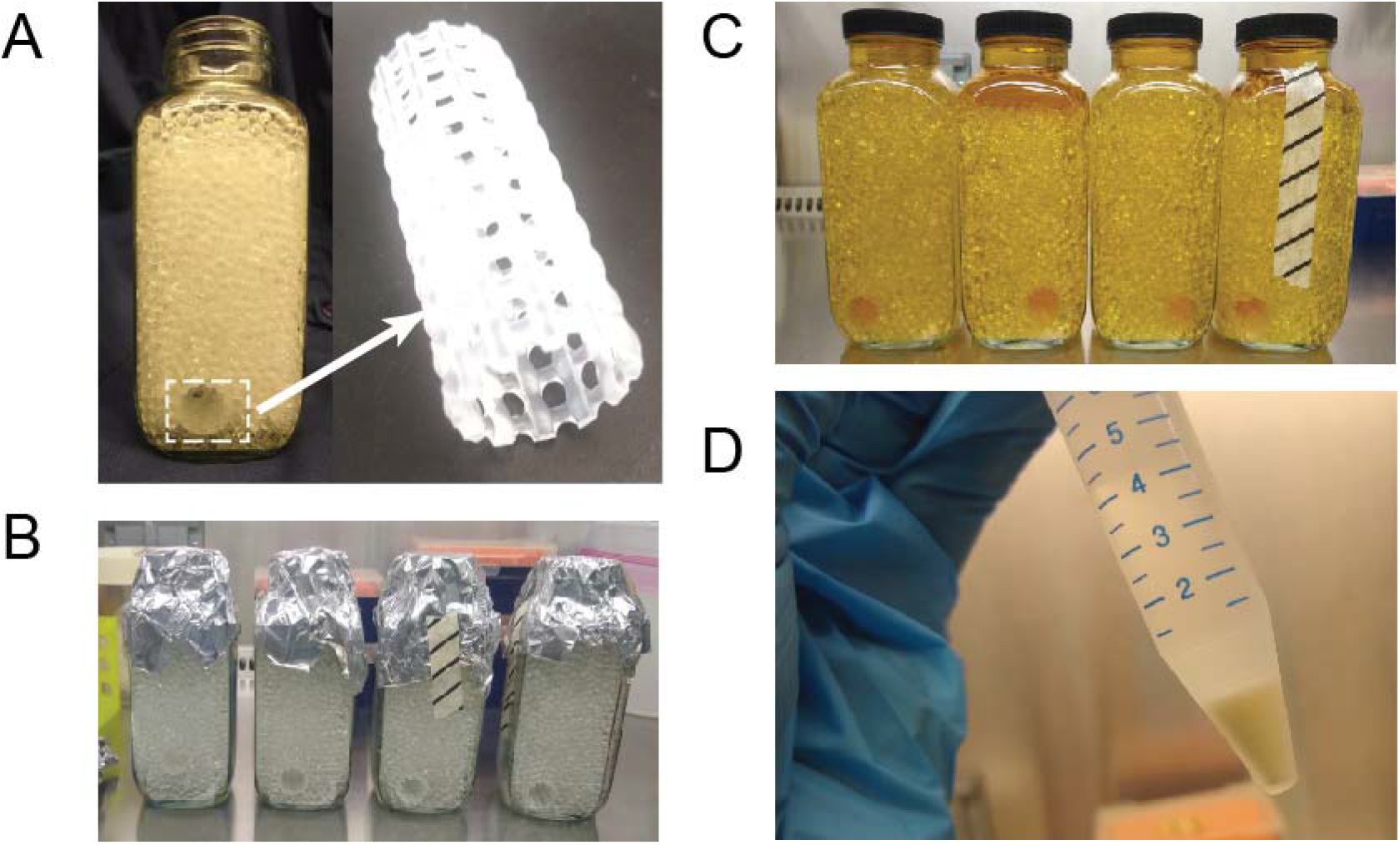
The bead/bottle system. **A. Left:** Square wide-mouth 480 mL French bottle fitted with a polypropylene mesh tube window and filled with 5 mm glass beads. **Right:** Closeup of polypropylene mesh tube “window”. **B.** View of bead-filled bottles after autoclaving. **C.** Inoculated bottles just before moving into incubator. **D.** Harvested Giardia pellet (about 10^9^ cells).

To sterilize this setup, the tops of the bead-filled bottles are covered in aluminum foil and autoclaved. The lids are autoclaved separately, because if autoclaved on the bottle, a seal can form while under vacuum and the hot polypropylene lid can significantly deform. To start the large-scale growth, media inoculated with about 3,000-5,000 cells per mL is added (about 225 mL per bottle) in a biosafety cabinet. To harvest, spent media is aspirated, detaching media (either pre-chilled or formononetin media) is added, and if chilling is used rather than formononetin, the bottles immersed in an ice/water slurry for 20 minutes. After mixing by repeated inversion, the solution of detached trophozoites is removed with a 25 mL disposable pipet. Note that an additional rinse step improves the yield only slightly.

The system features an extremely high surface-to-volume ratio – indeed, larger beads could be used – and so the yield depends on the volume and richness of the media. With normal media, late-log yields are about 500 million trophozoites per bottle and can be increased about twofold if richer media is used (see below). Because the growth is limited by the media rather by the available surface area, the adherent cells will reach only 25-50% confluence even at late log stage.

### Efforts to promote more vigorous growth with media enhancements

*Giardia* trophozoites have long been known to use arginine as a preferred energy source; the yield from TYI-S-33 media was doubled if supplemented with 10 mM arginine in the presence of 50 mM glucose (Edwards *et al*., 1992). Accordingly, using the bead/bottle system, we tested the effect of supplementing media with 5-50 mM arginine. In our hands, the improvement in yield with 10-20 mM arginine is much smaller, on the order of 20%. It is possible that the discrepancy is due to the use of different strains (WBC6 vs Portland I) or to differences in media components.

We have also used the bead/bottle system to test the effects of supplementing TYI-S-33 media with compounds identified as depleted in spent media (Table 3). Somewhat disappointingly, the only useful supplement tested is arginine-HCl at 10-20 mM (50 mM is worse). However, increasing the concentration of FBS to 20% along with adding arginine to 20 mM did significantly improve the media, with yields about twice that seen with normal media.

**Table 3.**
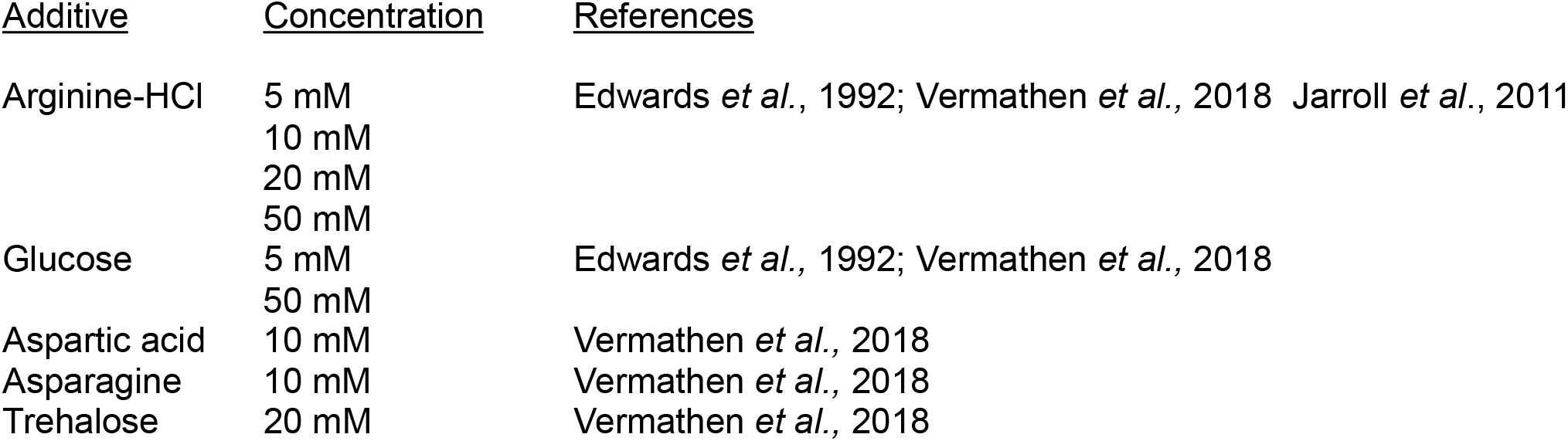
Media additives evaluated.

| <u>Additive</u> | <u>Concentration</u> | <u>References</u> |
| --- | --- | --- |
| Arginine-HCl | 5 mM | Edwards *et al.*, 1992; Vermathen *et al.*, 2018 Jarroll *et al.*, 2011 |
|  | 10 mM |  |
|  | 20 mM |  |
|  | 50 mM |  |
| Glucose | 5 mM | Edwards *et al.*, 1992; Vermathen *et al.*, 2018 |
|  | 50 mM |  |
| Aspartic acid | 10 mM | Vermathen *et al.*, 2018 |
| Asparagine | 10 mM | Vermathen *et al.*, 2018 |
| Trehalose | 20 mM | Vermathen *et al.*, 2018 |

### Troubleshooting

We have found several likely factors to troubleshoot if yields are unexpectedly low:

#### Media

The most crucial media components are the HI-FBS, the Biosate peptone (or, equivalently, the yeast extract and tryptone), the bile extract, and the cysteine-HCl. Problems with any of these will very likely result in poor growth. Ideally, new lots of the first three of these components should be tested, and if found suitable, larger quantities can then be purchased for the sake of consistency. Be sure to stir the media for 20-30 minutes before filtering.

#### Inoculum

Small batches of trophozoites should ideally be passaged at around 70-80% confluence, and allowing the trophozoites to become overgrown decreases yield at larger scales.

#### Cleaning

Before first use, all glassware should be soaked in dilute acid overnight and then thoroughly rinsed with MilliQ water. After each growth, all surfaces that have been in contact with *Giardia* must be disinfected and then rinsed extensively with water. If yields begin to drop over several growths, it may be necessary to clean the beads with a detergent such as MP Biomedical 7X detergent (Fisher ICN7667093), followed by extensive washing. A large, fritted glass funnel with a vacuum sidearm flask is very useful for accelerating the washing steps.

## Summary

As part of our efforts to improve methods for growing large scale preparations of *G. lamblia*, we have explored and optimized two new culture systems: the beehive and the bead/bottle. Compared to growth in standard 10 cm dishes, the bead/bottle method has additional benefits of using less media (and thus less FBS, the most expensive component of the system), taking up less room in incubators, and being more robust against accidental spills. In our hands, due to the ease of washing and of monitoring growth, the bead/bottle approach has proved invaluable in reliably increasing yields over traditional growth methods. In addition, we have modified and streamlined media preparation methods and identified common issues that can impair *G. lamblia* trophozoite growth. Together, we have developed a streamlined method for growing large preparations of *G. lamblia*, thus opening the door to structural biology, biophysical, and biochemical studies.

## Acknowledgments

The authors thank current and former Kieft and Rissland Lab members for thoughtful discussions and technical assistance. This work was supported by NIH grants R35GM128680 (OSR), R21AI149210 (BW & OSR) and R35GM118070 (JSK).

## Notes

### Competing Interest Statement

The authors have declared no competing interest.

